# Combinations of slow translating codon clusters can increase mRNA half-life

**DOI:** 10.1101/490995

**Authors:** Ajeet K. Sharma, Edward P. O’Brien

## Abstract

The introduction of a single, non-optimal codon cluster into a transcript was found to cause ribosomes to queue upstream of it and decrease the transcript’s half-life through the action of the dead box protein Dhh1p, which interacts with stalled ribosomes and promotes transcript degradation. Naturally occurring transcripts often contain multiple slow codon clusters, but the influence of these combinations of clusters on a transcript’s half-life is unknown. Using a kinetic model that describes translation and mRNA degradation, we find that the introduction of a second slow codon cluster into a transcript near the 5′-end can have the opposite effect than that of the first cluster by increasing the mRNA’s half-life. We experimentally validate this finding by showing that *S. cerevisiae* transcripts that only have a slow codon cluster towards the 3′-end of the coding sequence have shorter half-lives than transcripts with non-optimal clusters at both 3′ and 5′ -ends. The origin of this increase in half-life arises from the 5′-end cluster suppressing the formation of ribosome queues upstream of the second cluster, thereby decreasing the opportunities for translation-dependent degradation machinery, such as the Dhh1p protein, to interact with stalled ribosomes. We also find in the model that in the presence of two slow codon clusters the cluster closest to 5′-end is the primary determinant of mRNA half-life. These results identify two of the rules governing the influence of slow codon clusters on mRNA half-life, demonstrate that multiple slow codon clusters can have synergistic effects, and indicate that codon usage bias can play a more nuanced role in controlling cellular protein levels than previously thought.

## Introduction

The copy number of messenger RNA (mRNA) molecules *in vivo* is one of the primary determinants of cellular protein levels (1–3), and depends upon the rate of mRNA synthesis and degradation (1). Despite there being a small number of mechanisms by which mRNA molecules are degraded (4, 5) there is a wide variation in mRNA half-lives from a few minutes to over an hour in *S. cerevisiae* (6–9). Transcripts that code for ribosomal proteins, for example, tend to have longer half-lives as compared to the other transcripts (8). Such observations indicate that the nucleotide sequence of mRNA molecules are responsible for their differential degradation rates. Indeed, nucleotide sequence and structural features of the 5′ and 3′ untranslated regions were found to influence mRNA lifetime some two decades ago (10, 11). More recently, the nucleotide sequence within the coding region has also been shown to regulate mRNA degradation rates through the action of a protein that monitors ribosome traffic on transcripts (6, 12, 13). Thus, a range of sequence features can influence mRNA half-lives (5).

Within coding regions, enrichment of non-optimal codons (*i.e*., codons with relatively smaller cognate-tRNA concentrations) leads to faster degradation of the mRNA (6). In these cases, a translation-dependent degradation machinery recognizes and interacts with ribosomes slowed down by traffic-jams (*i.e*., queued ribosomes stacked next to one another on the transcript) caused by non-optimal codons that are translated more slowly than optimal codons (12). This interaction of translation-dependent degradation machinery with stalled ribosomes promotes the degradation of these mRNAs. Translation-dependent degradation can occur via multiple pathways that include the physical interaction of Dhh1p protein with stalled ribosomes (12) and the deadenylation of mRNA molecules with Ccr4-Not complex (14). The position of non-optimal codons within a transcript also affects mRNA half-life, with slow translating codon clusters located further from the start codon resulting in shorter half-lives (12). Evidence indicates this trend arises because longer ribosome queues can form when the non-optimal codon cluster is further from the start codon, giving more opportunities for the translation-dependent degradation machinery to engage stalled ribosomes. Translation-dependent degradation is thought to be beneficial for efficient global protein synthesis by avoiding the *sequestration* of a large number of ribosomes in non-productive traffic-jams (15).

To date, the effect of only a single slow codon cluster on an mRNA’s half-life has been studied (12). Yet in *S. cerevisiae* as many as 78% of transcripts contain two or more slow codon clusters, and theoretical studies have shown that multiple slow codon clusters can modulate the pattern of ribosome traffic-jams in complicated ways (16–18). Therefore, a richer set of mRNA lifetime behaviors might occur for many transcripts. In this study, we use kinetic modeling of translation and mRNA degradation to identify the scenarios and rules governing mRNA half-lives as two slow codon clusters are moved to different positions along a transcript’s coding sequence. We verify the simulation predictions against transcript half-lives that have been experimentally measured in *S. cerevisiae*.

## Methods

### Model for protein synthesis and mRNA degradation

We developed a computational model to study the effect of translation kinetics on mRNA degradation. In this model, a ribosome initiates translation with rate *α* when the first six codon positions of the transcript (19, 20) are not covered by another ribosome. An elongating ribosome nascent-chain complexes can then populate two distinct states depending upon the occupancy of the ribosome’s A-site. In state 1, the ribosome’s A-site is not occupied by an aminoacyl tRNA (aa-tRNA) while in state 2 it is (Fig. 1). A transition from the state 1 to 2 consists of the binding of a cognate aa-tRNA molecule and the covalent incorporation of the amino acid into the nascent chain. This step occurs with rate *ω*_a_(j) and is primarily determined by the cellular concentration of cognate and near-cognate tRNA molecules for the codon at the j^th^ position of the transcript. The ribosome-nascent chain complex is then translocated to the next codon position (j + 1) in state 1. This step occurs with rate *ω*_t_ when no ribosome is present *ℓ* codon positions downstream to the translocating ribosome, where *ℓ* is the number of consecutive codon positions covered by a ribosome. *ω*_’_ is assumed to be constant at all codon positions (21). These two steps elongate the nascent-chain by one amino acid and translocate the ribosome to the next codon position. Protein synthesis is terminated with rate *β* when the ribosome’s A-site arrives at the stop codon of the transcript resulting in the release of a fully synthesized protein.

**Figure 1:**
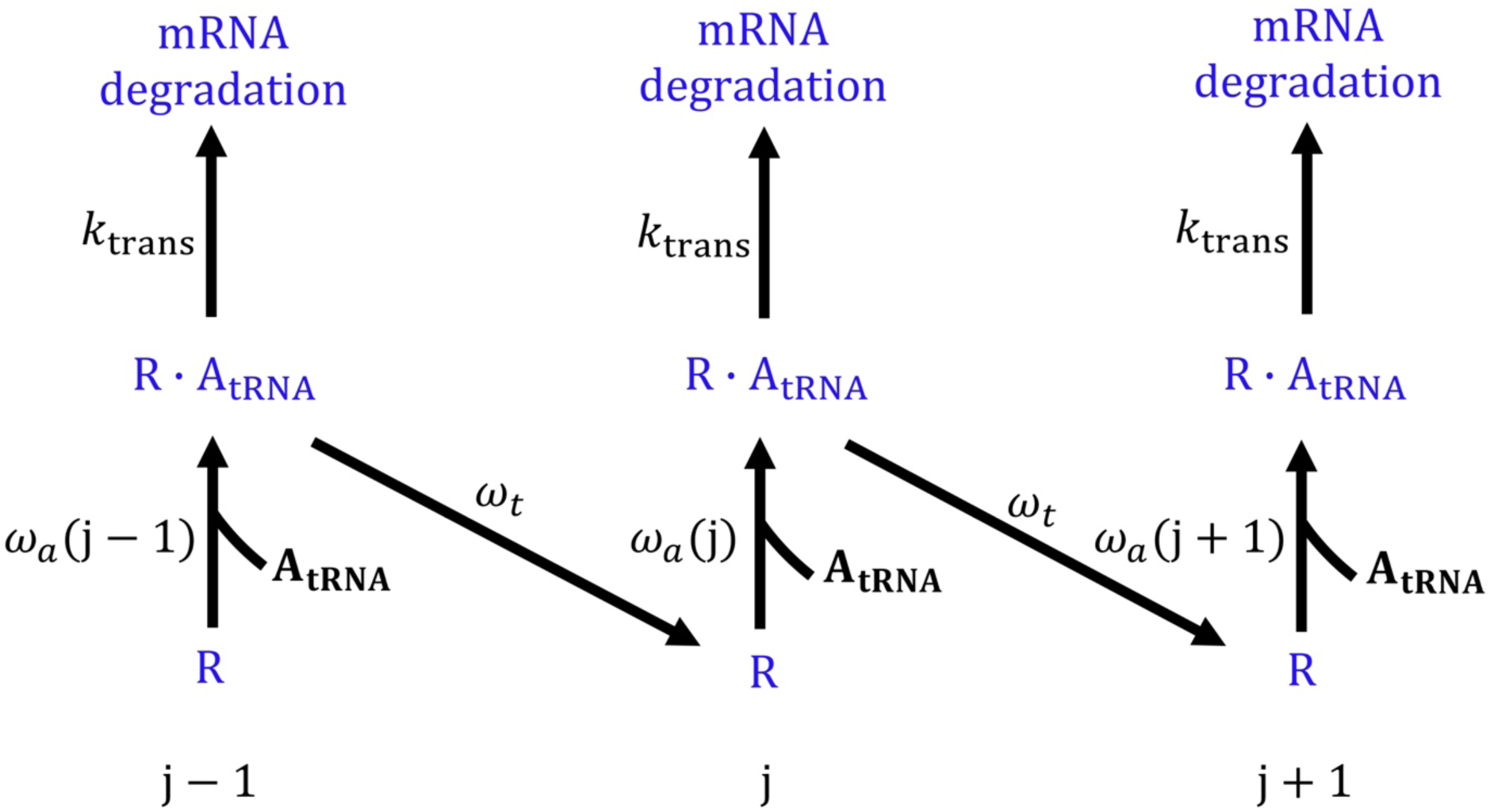
Kinetic scheme for protein synthesis and mRNA degradation. The binding of aa-tRNA (A_tRNA_) molecule to state 1 (R) of a ribosome-nascent chain complex at codon position j results in the transition to state 2 (R · A_tRNA_) at the rate *ω*_*a*_(j). An interaction between translation-dependent degradation machinery and state 2 of a ribosome-nascent chain complex leads to the degradation of mRNA molecule with the rate *k*_trans_. Alternatively, the ribosome nascent-chain complex can move to the next codon position in the state 1 with the rate *ω*_*t*_.

An mRNA molecule in our model is degraded by two independent mechanisms. First, it can be degraded by non-translation-dependent degradation mechanisms with rate *k*_mRNA_. These mechanisms include degradation through nonsense-mediated, no-go and non-stop decays (4). For simplicity, we assumed that the non-translation-dependent degradation pathways are independent of the coding sequence of the transcript. Second, in the reaction scheme shown in Fig. 1, the translation-dependent degradation machinery identifies state 2 of the ribosome nascent-chain complex and degrades the transcript with rate *k*_trans_. The assumption that translation-dependent degradation machinery can only interact to state 2 is motivated by two thought experiments. Consider the following two alternative scenarios. First, assume translation-dependent degradation machinery only interacts with state 1 (*i.e*., an empty A-site). In this case, the location of the slow codon cluster within a transcript would not affect the mRNA half-life as the traffic-jams on a transcript do not affect a ribosome’s dwell time in state 1. This contradicts the experimental observation in Ref. (12) that positioning the slow codon cluster further from start codon results in shorter mRNA half-lives. Second, assume translation-dependent degradation machinery interacts with both states 1 and 2. In this scenario the mRNA half-life will anti-correlate with transcript length as the average number of ribosomes on a transcript must increase with coding sequence length. However, plotting mRNA half-lives reported in Ref. (6) against coding sequence length does not exhibit this behavior (Fig. S1). Thus, the experimental observations are consistent with translation-dependent degradation machinery only interacting to state 2 (*i.e*., ribosomes in which a tRNA molecule in occupies the A-site).

We simulated protein synthesis and mRNA degradation using Gillespie’s algorithm (22); details are provided in the supporting information (SI). We start our simulations at *t* = 0 and then measure the time at which the mRNA is degraded by either of the two mechanisms of mRNA degradation. We repeated this procedure for 2,500 copies of each transcript examined and measured their turnover times. We then used these turnover times to calculate the decay of a transcript’s population as a function of time. The time when the transcript population is half its original value is the half-life of the transcript (8, 9).

### Parameters used in simulation model

In this model, codon translation times estimated by Fluitt and Viljoen are used (23) by breaking them into two parts. The first part consists of the time taken to bind aa-tRNA and incorporate an amino acid residue into the growing nascent-chain (*i.e*., transition from state 1 to 2) whereas the second part is the time spent translocating the ribosome to the next codon position (*i.e*., the transition from state 2 at codon position *j* to state 1 at codon position *j* + 1) (Fig. 1). We assumed that the ribosome translocation rate *ω*_t_ is 35 *s*^-1^ for all codon positions (21) whereas *ω*_*a*_(j) was calculated by subtracting the ribosome translocation time (*i.e*., the inverse of *ω*_*t*_) from the Fluitt-Viljoen codon translation times reported in Ref. (23). In the results presented we vary the translation-initiation rate α from 0.1 to 0.3 *s*^-1^ (21, 24, 25). We set *k*_mRNA_and *k*_trans_to 10^-4^ and 5×10^-5^ *s*^-1^, respectively, as these rate parameters give the PGK1 mRNA half-lives similar to the ones measured in Ref. (12). We set *β* and *ℓ* to 35 *s*^-1^ and 10, respectively (19, 21, 26).

### Construction of PGK1 variants

We constructed *in silico* a total of fifty-five variants of PGK1 transcripts. As in the experiments (12), ten variants were constructed by inserting a single cluster of ten non-optimal codons (TTA ATA GCG CGG CGG CGG CGG CGG GCG ACG) at codon positions 10, 20, …, 90 and 100% of the coding sequence length from the start codon in the wild-type PGK1 transcript. The remaining forty-five variants were constructed by then inserting a second cluster at different locations in the PGK1 transcript.

### Selection of three *S. cerevisiae* genes and the construction of their synonymous variants

We selected YDR158W, YPL061W and YLR109W *S. cerevisiae* transcripts to study the effects of multiple slow codon clusters on mRNA half-life. These transcripts have a relatively higher translation-initiation rate (25) as well as higher-than-average fractions of optimal codons that makes them more likely to have significant ribosome-traffic jams upon introducing non-optimal codons. We created four synonymous variants of each of these transcripts by replacing optimal codons with non-optimal ones (Table S1 and supporting file 1). To do this we scanned their coding sequences and found three positions where eight consecutive optimal codons can be replaced with non-optimal synonymous codons. Then two variants were constructed by replacing one of the three positions with non-optimal synonymous codons whereas the other two were constructed by replacing two of the three positions with non-optimal synonymous codons. The nucleotide sequences of those synonymous variants are provided in the Supporting file 1. While creating these synonymous variants, we considered a codon to be non-optimal if its tAI score is less than 0.47, otherwise the codon is an optimal codon (6, 27).

### Calculation of the average mRNA half-life using *in vivo* half-life data

We calculated the average half-lives for different sets of *S. cerevisiae* transcripts using the *in vivo* half-life data reported in Ref. (6). These sets of *S. cerevisiae* transcripts were constructed on the basis of whether slow codon clusters were present in the first and the last 20% of the coding sequences (*i.e*., towards the 5′ and 3′-end, respectively) of the transcripts (see Results). In this analysis, a slow codon cluster is defined as a stretch of six consecutive non-optimal codons in a transcript. We also varied the definition of a slow codon cluster by altering the size of the stretch of successive non-optimal codons from six to ten. The absence of a slow codon cluster in this analysis is identified as the absence of six or longer consecutive non-optimal codons. Note well, we did not vary the threshold of six non-optimal codons for the absence of a slow codon cluster while varying the definition of slow codon cluster. Moreover, to maintain sufficient sample sizes, the presence and absence of slow codon clusters in the middle 60% of the coding sequence was not considered.

## Results

### mRNA half-life decreases as a slow codon cluster is positioned further from the start codon

To test the accuracy of our model, which predicts mRNA half-lives by accounting for ribosome movement on transcripts and the action of translation-dependent degradation, we examined whether it can reproduce the experimentally observed trend that mRNA half-life decreases when a single slow codon cluster, consisting of ten non-optimal codons, moves away from the start codon (12). Specifically, the half-life of transcripts from gene PGK1 in *S. cerevisiae* decreases when this slow codon cluster is inserted 21 codons from the start codon and then moved to codon positions starting 104, 208, 263 and 321 codons from the start codon (12). We simulated protein synthesis and mRNA degradation for 2,500 copies of each of these five transcripts and calculated their half-lives (see Methods). Consistent with experiment (12), we find a decrease in mRNA half-life as the cluster of non-optimal codon is positioned further from the start codon (Fig. 2(A)). Since the initiation rate for the PGK1 transcript is unknown, we tested a range of biologically realistic initiation rates and find that the trend is robust (Fig. 2(A)). These results indicate the model can accurately describe the effect of slow codon clusters on mRNA half-life.

**Figure 2:**
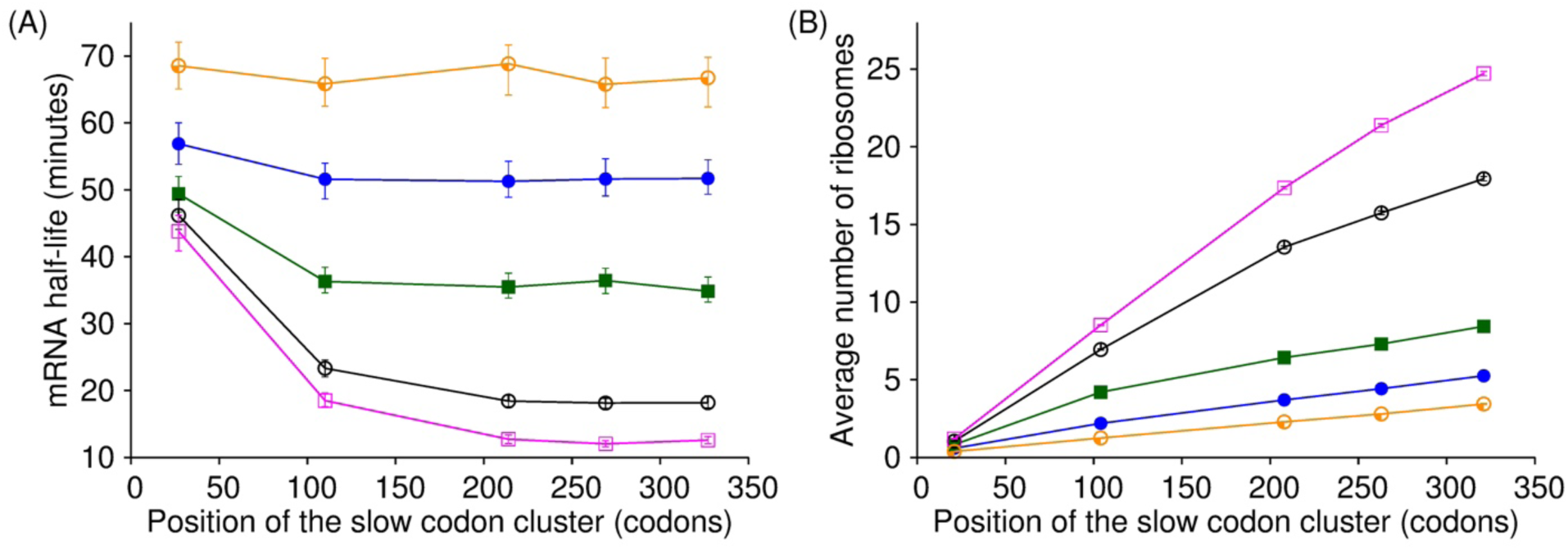
Half-life of *S. cerevisiae* PGK1 transcript decreases as a function of the position of slow codon cluster. PGK1 mRNA half-life (**A**) and the average number of ribosomes before the slow codon cluster (**B**) are plotted as a function of the position the cluster in the wild-type PGK1 transcript for *α* = 0.10, 0.15, 0.20, 0.25 and 0.30 *s*^-1^ in orange, blue, green, black and pink, respectively. Error bars in (**A**) and (**B**) are the 95% confidence interval which were calculated using 10,000 bootstrap cycles. Error bars in (**B**) are smaller than the symbol. Solid lines are to guide the eye.

We find that the average number of ribosomes queued upstream of the slow codon cluster increases when the cluster is further from the start codon (Fig. 2(B)), which supports Coller and coworker’s (12) molecular interpretation that as ribosome queues get longer the half-life decreases. The reason the queues are shorter when the slow codon cluster is nearer the start codon is that ribosome traffic-jams cannot backup beyond the translation initiation site, where ribosomes are assembled on the transcript. Additionally, the results in Fig. 2(B) explain why the dependence of half-life on cluster position is not as strong when the initiation rate decreases (Fig. 2(A)). As the initiation rate decreases the average ribosome density and queue length decreases, giving translation-dependent degradation machinery less opportunity to interact with stalled ribosomes and degrade the transcript.

### The first cluster is the primary determinant of mRNA half-lives

To understand the influence of multiple slow codon clusters we simulated how two clusters, located at different positions from the start codon, effect the PGK1 transcript’s half-life relative to PGK1 constructs containing one or zero clusters. A fundamental question in this two-cluster case is what the contribution of each cluster is to the transcripts half-life. To quantify this contribution we need to define several terms. The wild type transcript (which has zero slow-codon clusters) has half-life τ_0_; while the half-life of a transcript with one slow codon cluster, starting at codon position *i*, is τ_1,*i*_. Another cluster, starting at codon *k*, is then inserted downstream of the first cluster and that transcripts half-life is τ_2,*ik*_. In this case, the percent contribution of the first cluster to the half-life is

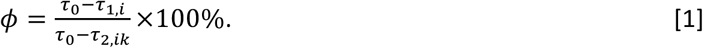

Note well, that for this analysis to be valid the location of the first cluster must be the same in the one-and two-cluster constructs, and the second cluster must be positioned downstream of the first. Defined in this way, ϕ is 0% when the first slow codon cluster does not contribute at all to the change in mRNA half-life upon going from the wild-type to two cluster construct (because τ_1,*i*_ will equal τ_0_); whereas all of the change in half-life comes from the first slow codon cluster when ϕ is 100% (because the numerator and denominator in Eq. 1 will be equal). We created 45 different constructs containing two slow codon clusters (see Methods). Plotting the parameter ϕ as a function of the position of the first and second cluster in the PGK1 transcript we find that in most cases the first cluster contributes more (ϕ > 50%) than the second cluster and in some cases it can be as much as 100% (Fig. 3(A)). Moreover, the effect of the first cluster becomes more prominent when it is positioned further from the start codon. We find similar results for a range of translation initiation rates (Figs. S2(A), (C), (E) and (G)). The reason the first cluster contributes more to translation-dependent degradation is that it acts as a bottleneck for ribosomes during translation, thereby reducing the average ribosome density and traffic-jams in the region between the first and the second cluster of the transcript (Figs. 3(B), S2(B), (D), (F) and (H)). This means translation-dependent degradation machinery can interact with more ribosomes positioned upstream of the first cluster than ribosomes translating codons between the first and the second cluster. Thus, the first codon cluster is the primary determinant of PGK1’s half-life when two clusters are present.

**Figure 3:**
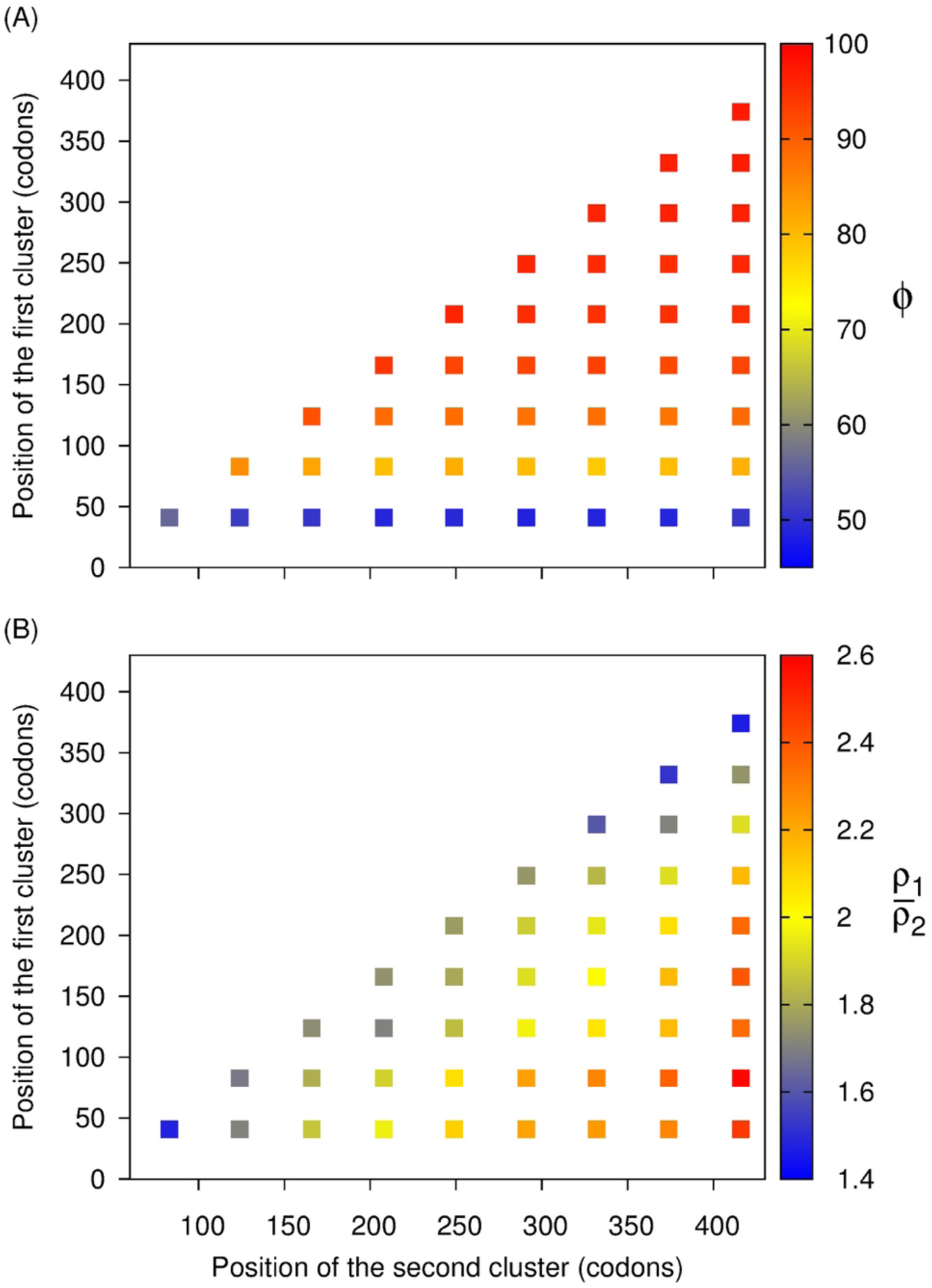
The first slow codon cluster in *S. cerevisiae* PGK1 controls its half-life. The heat map of the metric ϕ and the ratio of the average ribosome density before the first slow codon cluster to the average ribosome density between the first and second cluster 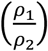 are plotted as function of the positions of the first and second cluster in the wild-type PGK1 transcript in (**A**) and (**B**), respectively.

To test whether this is a robust behavior in our model we simulated three other *S. cerevisiae* transcripts (YDR158W, YPL061W and YLR109W). We created three synonymous variants for each of these three transcripts by replacing eight consecutive optimal codons with non-optimal codons (see Methods), and calculated the mRNA half-lives for each of these synonymous variants using our simulation model. We find that ϕ varies up to 100% for these three *S. cerevisiae* transcripts, again indicating the majority contribution to translation-dependent degradation comes from the first slow codon cluster (Fig. 4).

**Figure 4:**
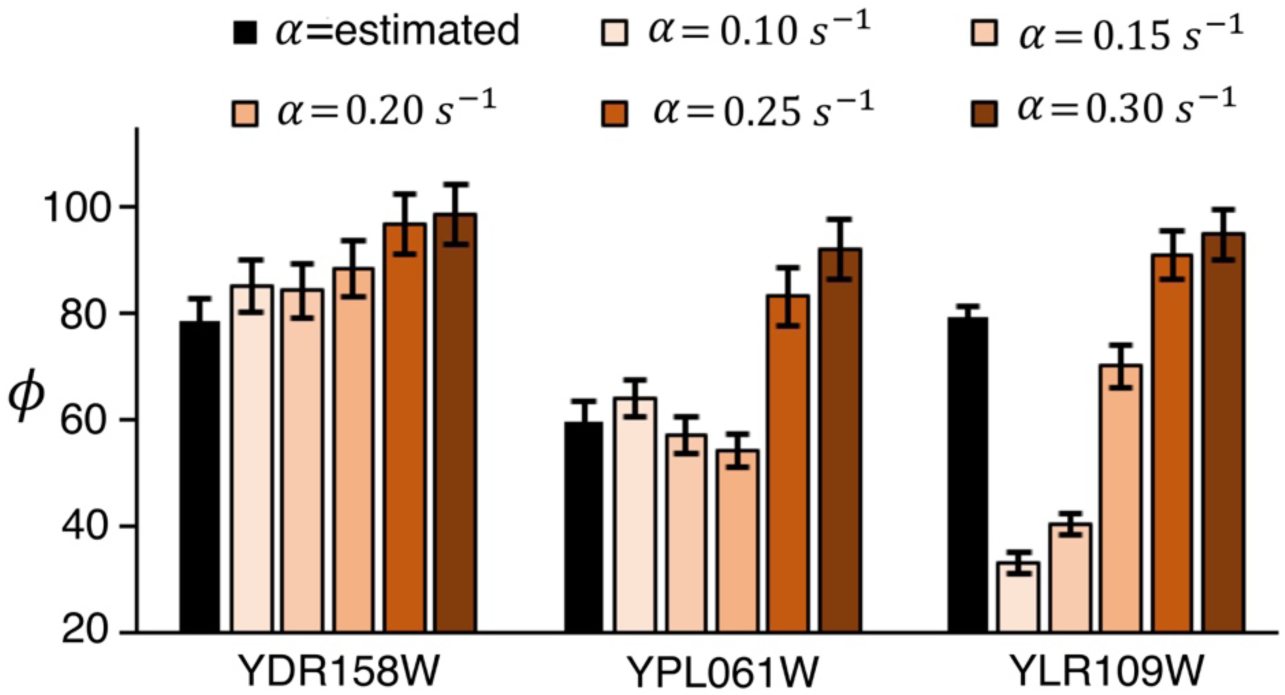
The first slow codon cluster in three *S. cerevisiae* transcripts controls their half-lives. The metric ϕ is plotted for six different translation-initiation rates using the half-lives of the wild-type and synonymous variants of YDR158W, YPL061W and YLR109W. The codon positions at which synonymous mutations are made in the wild-type transcripts to construct the synonymous variants are provided in the Table S1. The black bar represents the ϕ calculated from the mRNA half-lives measured in our simulations by using the translation-initiation rates reported in Ref. (25). Error bars are the 95% confidence interval which were calculated using 10,000 bootstrap cycles.

### Multiple slow codon clusters can increase mRNA half-life

Since the first slow cluster (*i.e*., nearest to the start codon) in a multi-cluster transcript determines the mRNA half-life (Fig. 3(A)), and the closer the first cluster is to the start codon the shorter the ribosome queues (Fig. 2(B)), we reasoned that introducing a new slow codon cluster near the start codon could increase the mRNA half-life by making the other existing clusters less relevant to mRNA degradation. This is opposite to the effect of a single slow codon cluster, which decreases mRNA half-life (12) (Fig. 2(A)). We tested this hypothesis by examining whether the simulated half-lives from our two-cluster constructs were greater than our single cluster constructs. Plotting the ratio τ_2_ by τ_1_, we find it is often greater than one in both PGK1 (Figs. 5 and S3) and the three *S. cerevisiae* constructs (Fig. 6) when the first cluster is closer to the start codon. This indicates that introducing a new slow codon cluster can increase an mRNA’s half-life. For two of the transcripts (YDR158W and YPL061W) the half-life is greater when two slow codon clusters are present than when none are present (Fig. S4), suggesting the increase in half life, for some transcripts, can be greater than the wild type.

**Figure 5:**
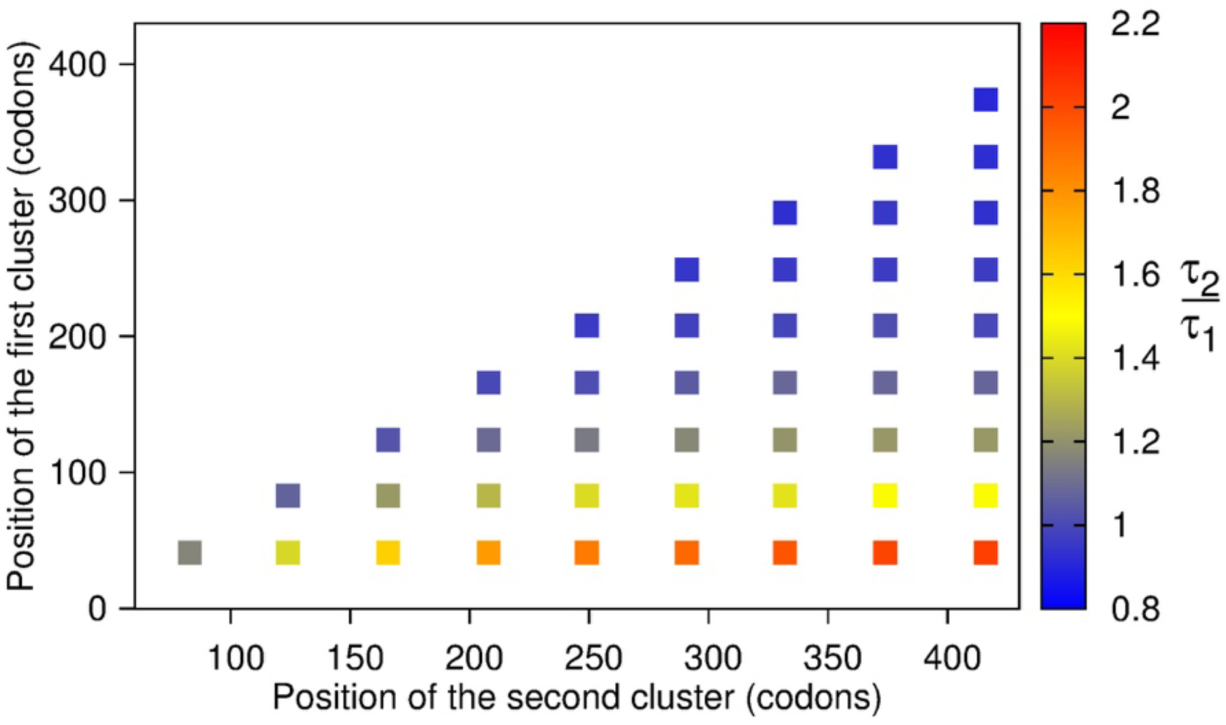
A slow codon cluster near the start codon in *S. cerevisiae* PGK1 transcript can increase its half-life. The heat map of 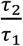 is plotted as a function of the position of the first and the second slow codon clusters in the wild-type PGK1 transcript.

**Figure 6:**
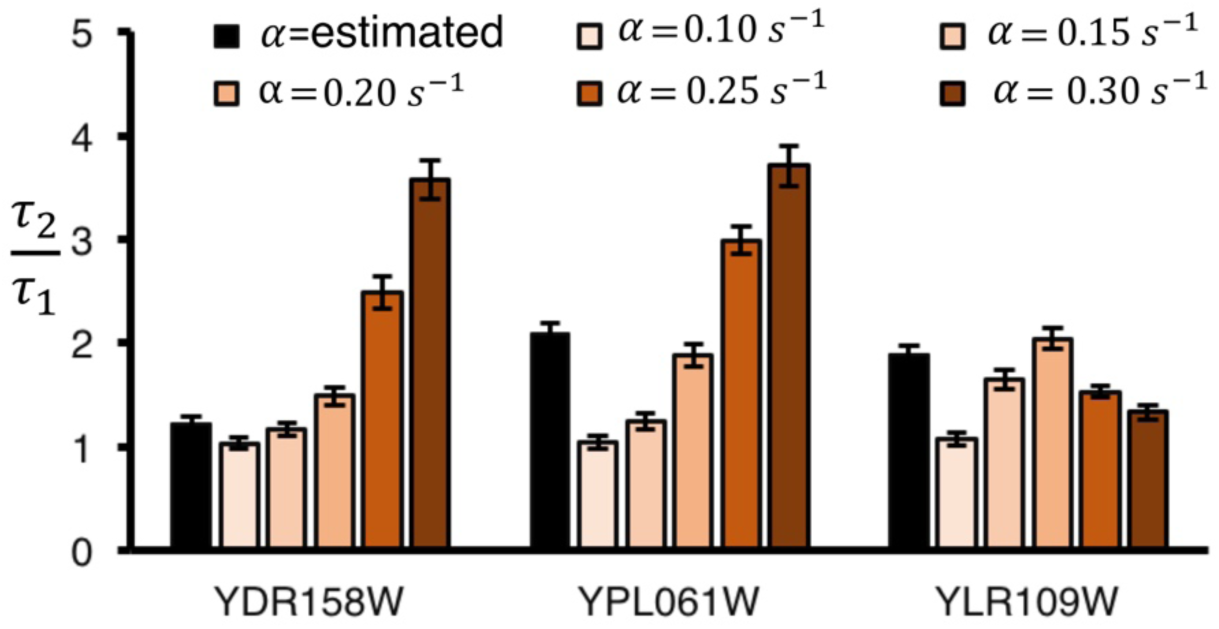
A slow codon cluster near the start codon increases the half-lives of *S. cerevisiae* transcripts. The ratio of the half-life of a transcript with two slow codon cluster to its synonymous variant with one slow codon cluster 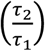 is plotted for six different translation-initiation rates. The black bar represents the ratio calculated from the mRNA half-lives measured in our simulations by using the translation-initiation rates reported in Ref. (25). The codon positions at which synonymous mutations are made in the wild-type transcripts to construct the synonymous variants are provided in the Table S1. Error bars are the 95% confidence interval which were calculated using 10,000 bootstrap cycles.

To experimentally test the predictions of our kinetic model we utilized *in vivo* mRNA half-life data measured in *S. cerevisiae* (6). Using these half-lives, we first reproduced the results of Ref. (12) at the transcriptome level showing that a slow codon cluster near the stop codon results in a decreased mRNA half-life (Supporting Results and Fig. S5). Next, we tested our prediction that the half-life can increase upon introduction of the second cluster near the 5′-end by examining whether the transcripts that had a slow codon cluster within the first 20% of the coding sequence (*i.e*., near the start codon) and the last 20% (*i.e*., near the stop codon) had longer half-lives than transcripts with slow codon clusters only in the last 20%. We indeed find that half-life is longer for the former (orange bars in Fig. 7) than the later (blue bars in Fig. 7). Thus, these experimental results support the kinetic model prediction – multiple slow codon clusters can increase mRNA half-life relative to a transcript with a single slow cluster.

**Figure 7:**
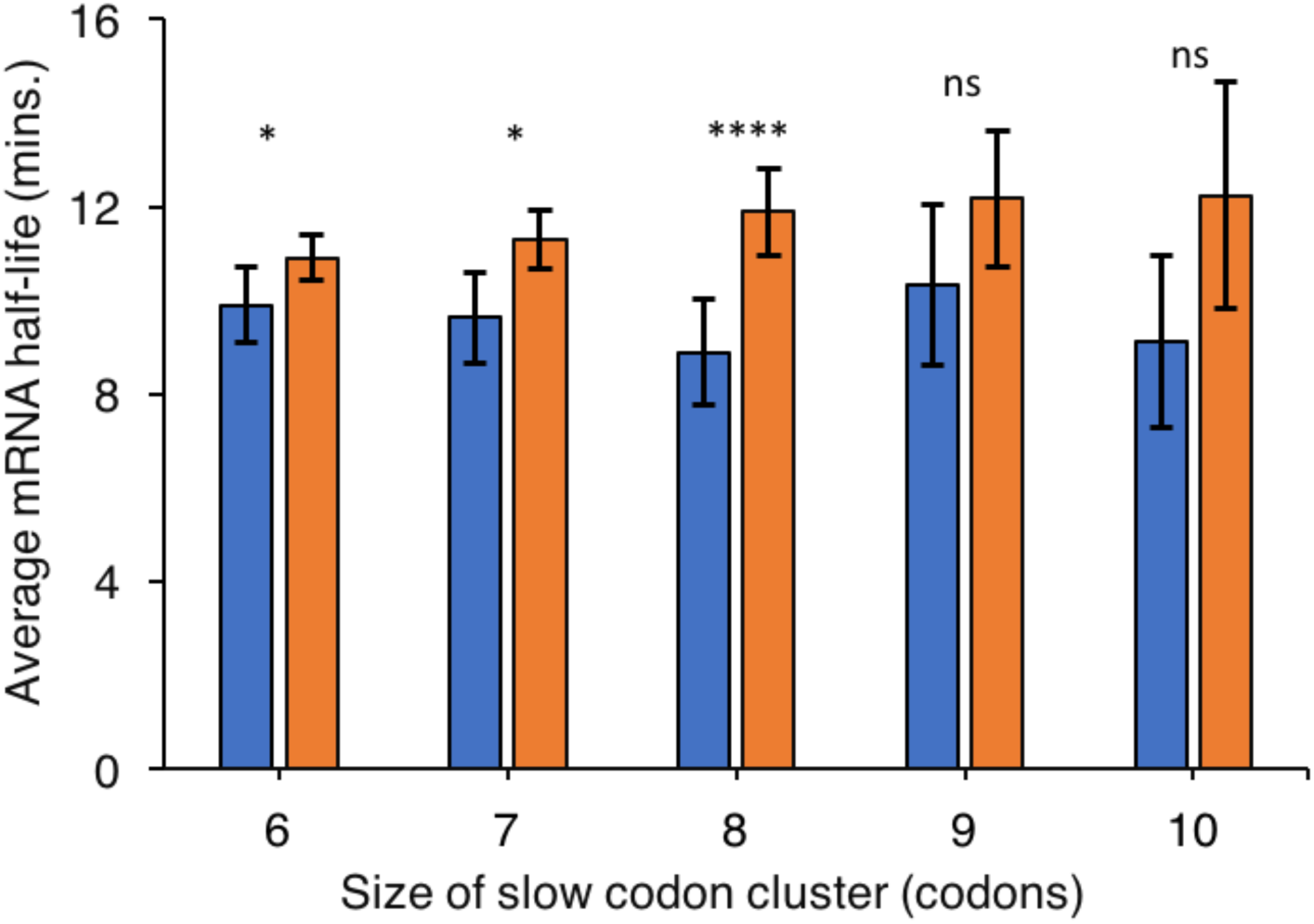
A slow codon cluster near the start codon increases mRNA half-life in *S. cerevisiae*. Average mRNA half-life in transcripts with slow codon clusters in the last 20% coding sequence and no slow codon cluster in the first 20% coding sequence (blue bars) is compared with transcripts with slow codon clusters in the first and the last 20% coding sequence (orange bars) for the varying definitions of slow codon cluster. The definition of a slow codon cluster is varied from six to ten consecutive non-optimal codons. Note well, the absence of a slow codon cluster is identified as the absence of six or more consecutive non-optimal codons for all bars. Error bars are the 95% confidence interval which were calculated using 10,000 bootstrap cycles. The number of transcripts used to calculate the average half-life for each bar is provided in the Table S2. (* denotes *p*-value < 0.05; **** denotes *p*-value < 10^-4^; ns denotes non-significant.)

## Discussion

In this study we have identified two rules governing the half-life of transcripts containing more than one non-optimal, slow-translating codon cluster: that the cluster closest to the 5′-end is the primary determinant of the half-life, and that the introduction of a second cluster into a transcript can increase a transcript’s half-life. We identified these rules from computer simulations of ribosomes translating transcripts and translation-dependent degradation, and then verified the second rule against the experimental data from *S. cerevisiae*. (The first rule could not be directly verified in this study as it requires new experiments involving synonymous substitutions within a single gene.)

The influence of these rules depends upon the position of the first slow codon cluster in the transcript. The first cluster has a greater impact when it is further from the start codon (Figs. 3 and S2) because it creates the greatest ribosome queues as compared to downstream cluster, and hence as it is positioned further from the start codon it can create even longer queues (Fig. 2(B)). Similarly, the increase in mRNA half-life due to the presence of two clusters becomes larger when the distance between them increases (Figs. 5 and S3). The reason for this is that the position of the first cluster is the primary determinant of mRNA half-life (Figs. 3 and S2), and the further this cluster is from the second cluster, the larger the decrease in ribosome traffic caused by the second cluster (Fig. 2(B)), and thus greater the increase in mRNA half-life.

A natural question is what, if any, biological benefit could be derived from these rules? We first note that slow codon clusters have been shown to aid in a variety of processes including co-translational protein folding (28, 29), protein targeting (30) and molecular recognition (31). But the introduction of slow codon clusters for one of these purposes could decrease the transcripts half-life. Therefore, the introduction of a second cluster, especially near the start codon, could offset this, maintaining the biological benefit of the original cluster with none of the negative effects on mRNA half-life. In fact, a study has found an enrichment of non-optimal codons in the first fifty codons of transcripts across the twenty-seven different organisms (32). In that study, it was proposed that this “ramp of slow codons” increases translation efficiency by reducing ribosome traffic-jams. Our results suggest that an additional benefit of such conservation of non-optimal codons is that it can increase mRNA half-lives.

We also found that increasing a transcript’s translation-initiation rate decreases its half-life because it creates more ribosome traffic jams allowing more translation-dependent degradation (Fig. 2(A)). This observation suggests that the presence of mRNA features that give rise to slower initiation rates in a transcript should positively correlate with longer mRNA half-lives. For example, more stable structure near the 5′ UTR decreases translation-initiation rates (24, 25, 33). Consistent with this reasoning a strong positive correlation has been reported between the free energy of mRNA folding near the 5′ UTR and mRNA half-life (34).

The mRNA half-life rules we have discovered are most applicable to transcripts where non-optimal codons can significantly modulate ribosome traffic-jams. Ribosome traffic on transcripts with low initiation rates, for example, are less likely to be affected by non-optimal codons due to the low ribosome density on those transcripts (Fig. 2), and less likely to exhibit variation in translation-dependent degradation rates. Therefore, the consequences of these rules are most likely to manifest themselves in highly expressed transcripts that tend to have higher translation-initiation rates (25). In summary, this work highlights the broader role played by non-optimal codons in regulating gene expression and expands our understanding of the molecular origin of differential degradation rates of transcripts.

## Supporting information

