## Supplementary material for "Combinations of slow translating codon clusters can increase mRNA half-life"

**Supporting methods.** We simulated the protein synthesis and mRNA degradation using Gillespie's algorithm (1). To do that first we define a parameter

$$R = \alpha + \sum_{j=2}^{N_c-1} \omega_a(j)\delta_1(j) + \sum_{j=2}^{N_c-1} \omega_a(j)\delta_2(j) + \beta + \sum_{j=2}^{N_c-1} \delta_2(j)\, k_{\mathrm{trans}} + k_{\mathrm{mRNA}} \qquad \text{[S1]}$$

where $\delta_{\mathrm{k}}(j) = 1$ when a ribosome occupies the j$^{th}$ codon position in the state $\mathrm{k}$, otherwise it is zero. The time interval between two successive transitions in the Gillespie's algorithm is exponentially distributed with a mean value of $\frac{1}{R}$. Therefore, we generate an exponentially distributed random number

$$\tau = \frac{1}{R} \ln{(r_1)}, \qquad \text{[S2]}$$

that provides the time for the next transition in our simulations. $r_1$ in Eq. [S2] is a random number that is uniformly distributed between 0 and 1. We generated another uniform random number $r_2$ between 0 and $R$ to identify the next step in our simulations according to the algorithm provided in the Table S3.

### Supporting results.

**Slow codon clusters near the 3′-end of a transcripts results in shorter mRNA half-lives.** Coller and co-workers find that introducing a slow codon cluster near the 3′-end of the coding sequence decreases the half-life of the PGK1 transcript (2). We tested whether this is also true at the transcriptome level by comparing the average mRNA half-life of transcripts with slow codon clusters in the last 20% coding sequence and no slow codon cluster in the first 20% coding sequence with transcripts with no slow codon cluster in the first and last 20% coding sequence. We found that the average half-life is longer for the former (blue bars in Fig. S5) than the later (orange bars in Fig. S5). Thus, these results provide an experimental evidence that slow codon clusters near the 3′-end of transcripts result in shorter mRNA half-lives even at the transcriptome level.

**Table S1.** Codon positions where synonymous mutations are made in Figs. 5 and 6.

| *S. cerevisiae* transcript | Synonymous mutations in Fig. 5 (codon positions) | Synonymous mutations in Fig. 6 (codon positions) |
|---|---|---|
| YDR158W | Variant 1: 326-333<br>Variant 2: 326-333 and 350-357 | Variant 1: 350-357<br>Variant2: 14-21 and 350-357 |
| YPL061W | Variant 1: 466-473<br>Variant 2: 466-473 and 491-498 | Variant 1: 491-498<br>Variant 2: 14-31 and 491-498 |
| YLR109W | Variant 1: 129-136<br>Variant 2: 129-136 and 145-152 | Variant 1: 145-152<br>Variant 2: 17-24 and 145-152 |

**Table S2.** List of the number of *S. cerevisiae* transcripts used in Fig. 7 for calculating average mRNA half-life.

| Size of slow codon cluster | Number of transcripts used for calculating blue bars | Number of transcripts used for calculating orange bars |
|---|---|---|
| 6 | 519 | 1759 |
| 7 | 332 | 1059 |
| 8 | 203 | 544 |
| 9 | 113 | 244 |
| 10 | 69 | 97 |

**Table S3.** A list of the types of transitions and the conditions for those transitions in our simulations.

| Conditions | Transitions |
|---|---|
| $r_2 < \alpha$ and the first six codon positions of the transcripts are unoccupied | Translation-initiation |
| $\alpha \le r_2$, $\sum_{i=2}^{k-1} \omega_a(i)\delta_1(i) + \alpha \le r_2 < \sum_{i=2}^{k} \omega_a(i)\delta_1(i) + \alpha$ and a ribosome is at the $k^{th}$ codon position in state 1 | Transition of the ribosome nascent chain complex from state 1 to 2 at the $k^{th}$ codon position |
| $\sum_{i=2}^{N_c-1} \omega_a(i)\delta_1(i) + \sum_{i=2}^{k-1} \omega_a(i)\delta_2(i) + \alpha \le r_2 < \sum_{i=2}^{N_c-1} \omega_a(i)\delta_1(i) + \sum_{i=2}^{k} \omega_a(i)\delta_2(i) + \alpha, \quad k > N_c - 10$ and a ribosome is at the $k^{th}$ codon position in state 2 with no ribosome at $k + 10$. | Transition of the ribosome nascent chain complex from the state 2 of the $k^{th}$ codon position to the state 1 of $(k+1)^{th}$ codon position |
| $\sum_{i=2}^{N_c-1} \omega_a(i)\delta_1(i) + \sum_{i=2}^{k-1} \omega_a(i)\delta_2(i) + \alpha \le r_2 < \sum_{i=2}^{N_c-1} \omega_a(i)\delta_1(i) + \sum_{i=2}^{k} \omega_a(i)\delta_2(i) + \alpha,\ N_c - 10 \le k < N_c$ and a ribosome is at the $k^{th}$ codon position in state 2 | Transition of the ribosome nascent chain complex from the state 2 ($k^{th}$ codon position) to the state 1 ($(k+1)^{th}$ codon) |
| $\sum_{i=2}^{N_c-1} \omega_a(i)\delta_1(i) + \sum_{i=2}^{N_c-1} \omega_a(i)\delta_2(i) + \alpha \le r_2 < \sum_{i=2}^{N_c-1} \omega_a(i)\delta_1(i) + \sum_{i=2}^{N_{\square}-1} \omega_a(i)\delta_2(i) + \beta$ and a ribosome at the stop codon | Translation-termination |
| $\sum_{i=2}^{N_c-1} \omega_a(i)\delta_1(i) + \sum_{i=2}^{N_c-1} \omega_a(i)\delta_2(i) + \alpha + \beta \le r_2 < \sum_{i=2}^{N_c-1} \omega_a(i)\delta_1(i) + \sum_{i=2}^{N_c-1} \omega_a(i)\delta_2(i) + \sum_{j=2}^{N_c-1} \delta_2(j)\, k_{\text{trans}}$ | Translation dependent degradation of mRNA |
| $\sum_{i=2}^{N_c-1} \omega_a(i)\delta_1(i) + \sum_{i=2}^{N_c-1} \omega_a(i)\delta_2(i) + \alpha + \beta + \sum_{j=2}^{N_c-1} \delta_2(j)\, k_{\text{trans}} \le r_2 < \sum_{i=2}^{N_c-1} \omega_a(i)\delta_1(i) + \sum_{i=2}^{N_c-1} \omega_a(i)\delta_2(i) + \sum_{j=2}^{N_c-1} \delta_2(j)\, k_{\text{trans}} + k_{\text{mRNA}}$ | Non-translation dependent degradation of mRNA |

**Table S4.** List of the number of *S. cerevisiae* transcripts used in Fig. S5 for calculating average mRNA half-life.

| Size of slow codon cluster | Number of transcripts used for calculating blue bars | Number of transcripts used for calculating orange bars |
|---|---|---|
| 6 | 890 | 519 |
| 7 | 890 | 332 |
| 8 | 890 | 203 |
| 9 | 890 | 113 |
| 10 | 890 | 69 |

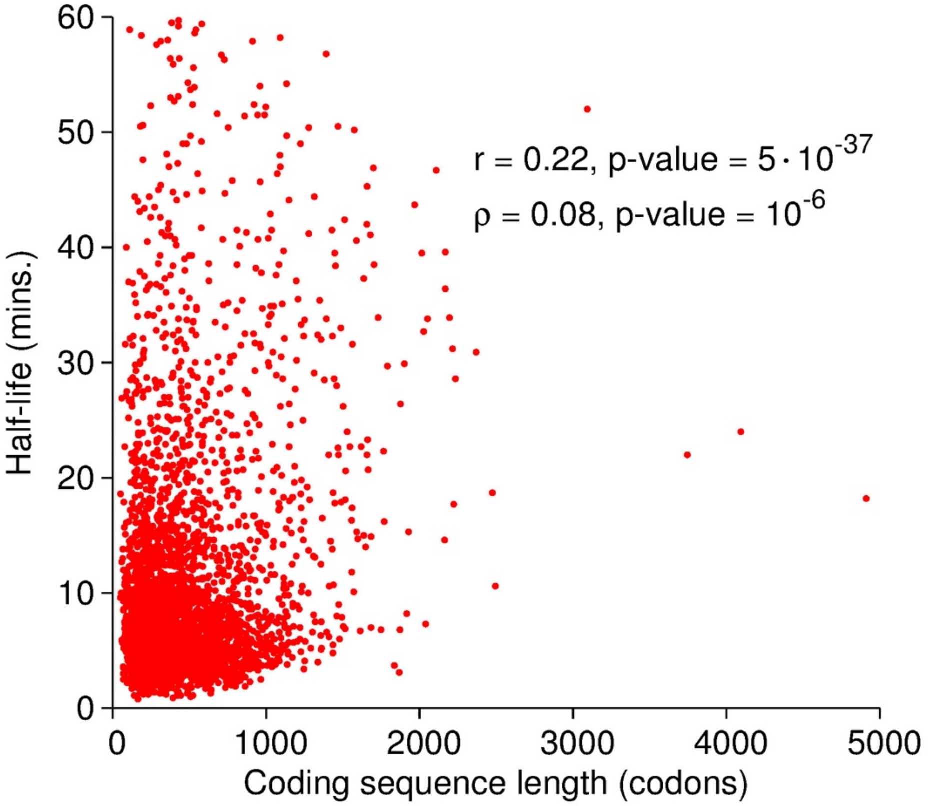


**Figure S1:** mRNA half-life correlates with coding sequence length in *S. cerevisiae*.

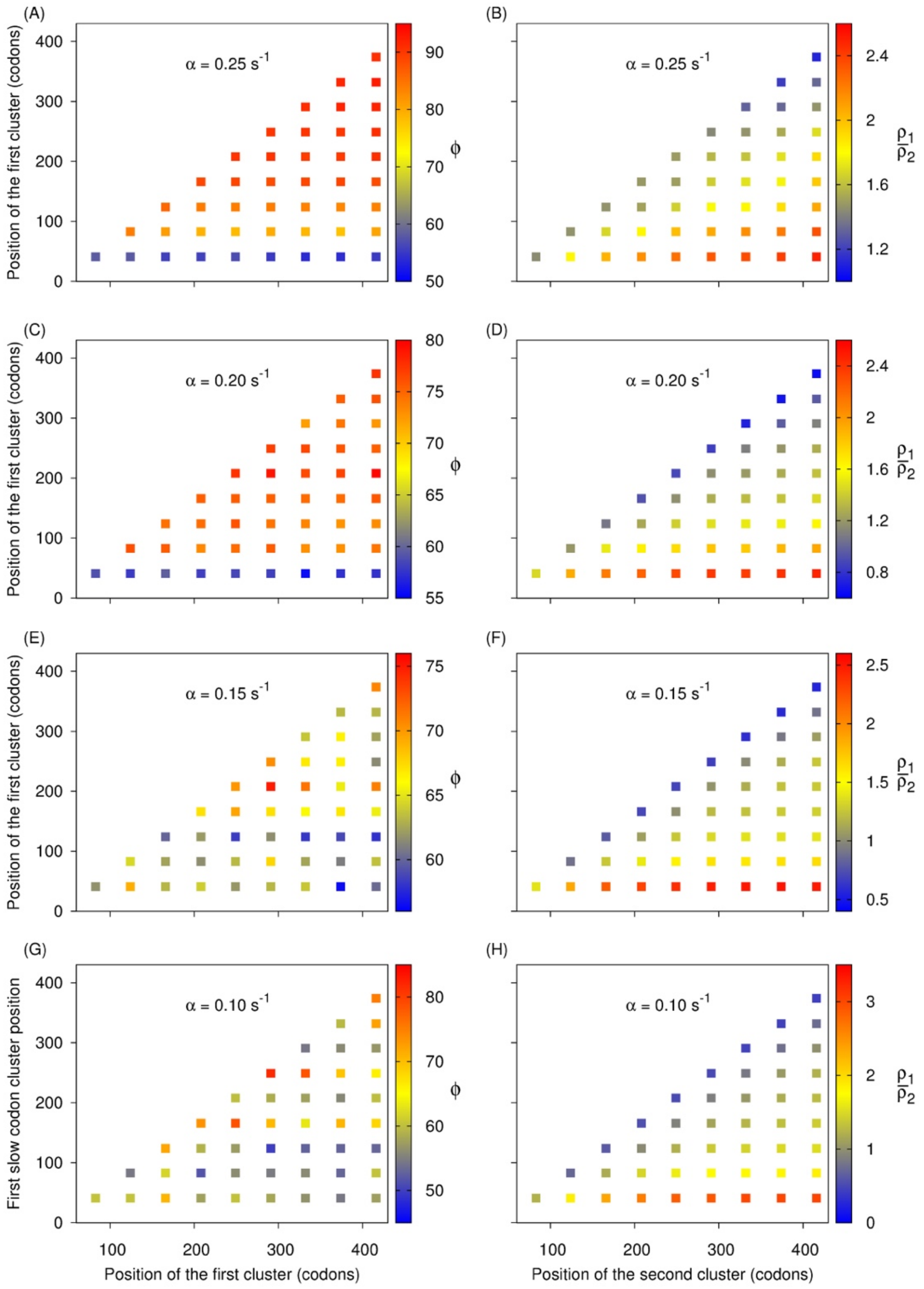


**Figure S2. The first slow codon cluster in *S. cerevisiae* PGK1 transcript controls its half-life.** Heat map of the metric $\phi$ ((**A**), (**C**), (**E**) and (**G**)) and the ratio of the average ribosome density before the first cluster to the average ribosome density between the first and second cluster $\left(\frac{\rho_1}{\rho_2}\right)$ ((**B**), (**D**), (**F**) and (**H**)) are plotted as a function of the position of the first and the second slow codon cluster in the wild-type PGK1 transcript. (**A**) and (**B**) are created by setting initiation rate $\alpha = 0.25\ s^{-1}$ in our simulations; (**C**) and (**D**) are created by setting initiation rate $\alpha = 0.20\ s^{-1}$ in our simulations; (**E**) and (**F**) are created by setting initiation rate $\alpha = 0.15\ s^{-1}$ in our simulations; and (**G**) and (**H**) are created by setting initiation rate $\alpha = 0.10\ s^{-1}$ in our simulations.

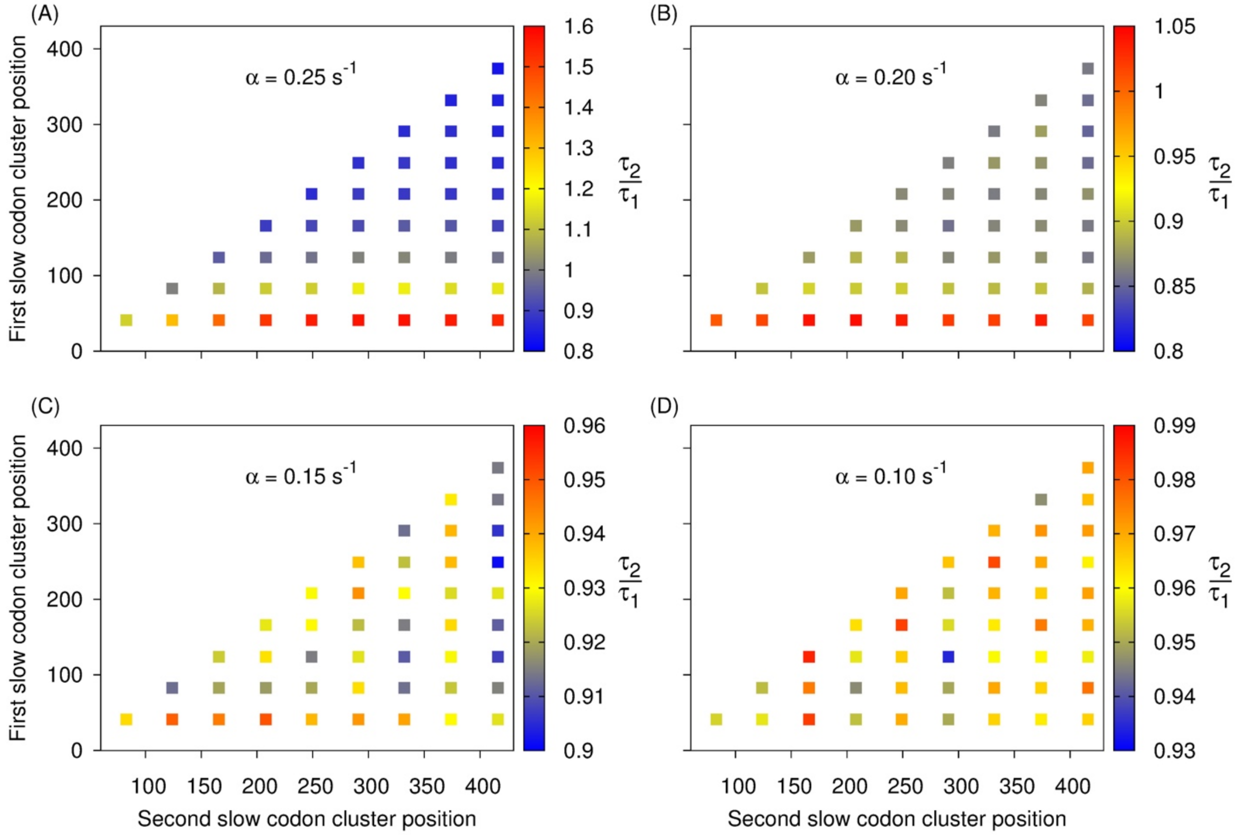


**Figure S3**: **A slow codon cluster near the start codon in *S. cerevisiae* PGK1 transcript can increase its half-life.** Heat map of the metric $\frac{\tau_2}{\tau_1}$ is plotted as a function of the position of the first and the second slow codon cluster in the wild-type PGK1 transcript in (**A**), (**B**), (**C**) and (**D**) using the initiation rates $0.25, 0.20, 0.15$ and $0.10\ s^{-1}$, respectively.

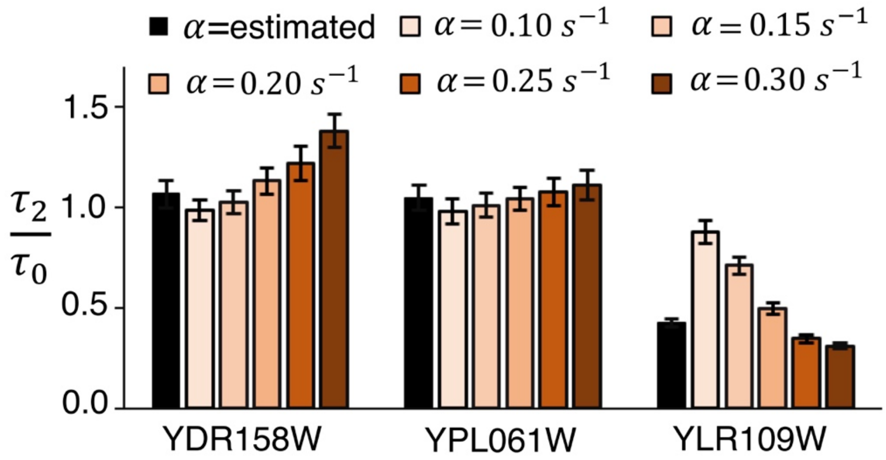


**Figure S4**: **Synonymous variants with two slow codon clusters can have longer half-lives than the wild-type *S. cerevisiae* transcripts.** The ratio of the half-life of transcript with two slow codon clusters to the half-life of the wild-type *S. cerevisiae* transcripts $\left(\frac{\tau_2}{\tau_0}\right)$ is plotted for six different translation-initiation rates. The black bar represents the ratio calculated from the mRNA half-lives measured in our simulations by using the translation-initiation rates reported in Ref. (3). Error bars are the 95% confidence interval which were calculated using 10,000 bootstrap cycles.

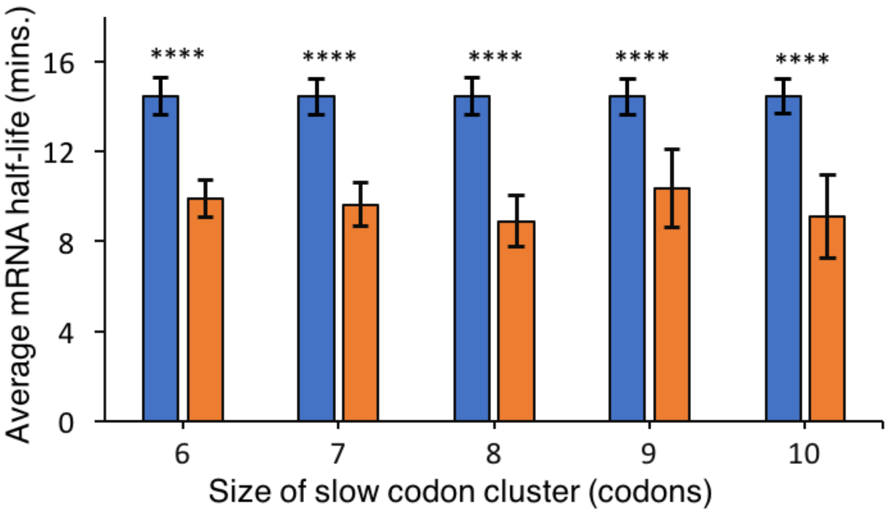


**Figure S5**: **A slow codon cluster near the stop codon results in shorter mRNA half-lives in *S. cerevisiae* transcripts.** Average mRNA half-life in transcripts with slow codon clusters in the last 20% coding sequence and no slow codon cluster in the first 20% coding sequence (orange bars) is compared with transcripts with no slow codon cluster in the first and last 20% coding sequence (blue bars) for the varying definitions of a slow codon cluster. The definition of a slow codon cluster is varied from six to ten consecutive non-optimal codons. Note well, the absence of a slow codon cluster is identified as the absence of six or more consecutive non-optimal codons in all bars. The number of transcripts used to calculate the average half-life for each bar is provided in the Table S4. (**** denotes $p$-value $< 10^{-4}$)
