## Supplementary material for "Combinations of slow translating codon clusters can increase mRNA half-life"

>double_cluster_1_3_YDR158W (used to calculate $\tau_2$)
ATGGCTGGAAAGAAAATTGCTGGTGTTTTGGGTGCTACTGGGTCGGTAGGGCAGAGGTTC
ATACTGTTGTTGGCAAATCACCCTCATTTCGAACTGAAAGTTCTTGGTGCCTCTTCTAGA
TCAGCTGGCAAGAAATACGTTGACGCTGTGAACTGGAAGCAAACCGATTTGCTACCGGAA
TCTGCTACCGATATTATTGTTTCCGAATGTAAATCTGAATTCTTTAAAGAGTGTGACATC
GTCTTTTCCGGATTGGATGCTGACTATGCTGGCGCTATCGAAAAGGAATTCATGGAAGCT
GGTATCGCCATTGTTTCCAATGCCAAGAATTATAGAAGAGAACAAGATGTGCCATTGATT
GTTCCTGTTGTCAATCCTGAGCATTTGGATATTGTAGCTCAAAAGCTTGACACCGCCAAG
GCTCAAGGTAAGCCAAGACCAGGGTTCATTATCTGTATTTCCAATTGTTCCACTGCAGGT
TTGGTTGCACCATTGAAGCCTTTGATTGAAAAATTCGGTCCTATTGATGCTTTGACCACT
ACTACTTTGCAAGCAATCTCAGGTGCTGGTTTCTCCCCAGGTGTACCAGGTATTGATATT
CTAGACAATATTATTCCATACATTGGTGGTGAAGAAGACAAGATGGAATGGGAGACCAAG
AAAATCTTGGCTCCATTAGCAGAAGACAAGACACACGTCAAACTATTGACTCCAGAAGAA
ATCAAAGTCTCTGCTCAATGTAACAGAGTCGCTGTTTCCGATGGGCACACCGAATGTATC
TCTTTGAGGTTCAAGAACAGACCTGCTCCATCCGTCGAGCAAGTCAAGACATGCCTAAAA
GAATACGTCTGCGATGCCTACAAATTAGGCTGTCATTCTGCTCCAAAGCAAACTATTCAT
GTTTTGGAACAACCAGACAGACCTCAACCAAGGTTGGACAGGAACAGAGACAGCGGTTAC
GGTGTTTCCGTTGGTAGAATCAGAGAAGACCCATTGTTAGATTTCAAAATGGTTGTCCTT
TCCCACAACACCATTATTGGTGCCGCTGGGTCGGGGGTACTTATAGCAGAGATCTTACTA
GCAAGAAACTTGATTTAA

>double_cluster2_3_YDR158W (used to calculate $\tau_{2,ik}$)
ATGGCTGGAAAGAAAATTGCTGGTGTTTTGGGTGCTACTGGTTCCGTTGGTCAACGTTTC
ATTCTGTTGTTGGCAAATCACCCTCATTTCGAACTGAAAGTTCTTGGTGCCTCTTCTAGA
TCAGCTGGCAAGAAATACGTTGACGCTGTGAACTGGAAGCAAACCGATTTGCTACCGGAA
TCTGCTACCGATATTATTGTTTCCGAATGTAAATCTGAATTCTTTAAAGAGTGTGACATC
GTCTTTTCCGGATTGGATGCTGACTATGCTGGCGCTATCGAAAAGGAATTCATGGAAGCT
GGTATCGCCATTGTTTCCAATGCCAAGAATTATAGAAGAGAACAAGATGTGCCATTGATT
GTTCCTGTTGTCAATCCTGAGCATTTGGATATTGTAGCTCAAAAGCTTGACACCGCCAAG
GCTCAAGGTAAGCCAAGACCAGGGTTCATTATCTGTATTTCCAATTGTTCCACTGCAGGT
TTGGTTGCACCATTGAAGCCTTTGATTGAAAAATTCGGTCCTATTGATGCTTTGACCACT
ACTACTTTGCAAGCAATCTCAGGTGCTGGTTTCTCCCCAGGTGTACCAGGTATTGATATT
CTAGACAATATTATTCCATACATTGGTGGTGAAGAAGACAAGATGGAATGGGAGACCAAG
AAAATCTTGGCTCCATTAGCAGAAGACAAGACACACGTCAAACTATTGACTCCAGAAGAA
ATCAAAGTCTCTGCTCAATGTAACAGAGTCGCTGTTTCCGATGGGCACACCGAATGTATC
TCTTTGAGGTTCAAGAACAGACCTGCTCCATCCGTCGAGCAAGTCAAGACATGCCTAAAA
GAATACGTCTGCGATGCCTACAAATTAGGCTGTCATTCTGCTCCAAAGCAAACTATTCAT
GTTTTGGAACAACCAGACAGACCTCAACCAAGGTTGGACAGGAACAGAGACAGCGGTTAC
GGTGTTTCCGTTGGTAGGATAAGGGAGGATCCCCTTCTTGATTTCAAAATGGTTGTCCTT
TCCCACAACACCATTATTGGTGCCGCTGGGTCGGGGGTACTTATAGCAGAGATCTTACTA
GCAAGAAACTTGATTTAA

>wildtype_YDR158W (used to calculate $\tau_0$)
ATGGCTGGAAAGAAAATTGCTGGTGTTTTGGGTGCTACTGGTTCCGTTGGTCAACGTTTC
ATTCTGTTGTTGGCAAATCACCCTCATTTCGAACTGAAAGTTCTTGGTGCCTCTTCTAGA
TCAGCTGGCAAGAAATACGTTGACGCTGTGAACTGGAAGCAAACCGATTTGCTACCGGAA
TCTGCTACCGATATTATTGTTTCCGAATGTAAATCTGAATTCTTTAAAGAGTGTGACATC
GTCTTTTCCGGATTGGATGCTGACTATGCTGGCGCTATCGAAAAGGAATTCATGGAAGCT
GGTATCGCCATTGTTTCCAATGCCAAGAATTATAGAAGAGAACAAGATGTGCCATTGATT
GTTCCTGTTGTCAATCCTGAGCATTTGGATATTGTAGCTCAAAAGCTTGACACCGCCAAG
GCTCAAGGTAAGCCAAGACCAGGGTTCATTATCTGTATTTCCAATTGTTCCACTGCAGGT
TTGGTTGCACCATTGAAGCCTTTGATTGAAAAATTCGGTCCTATTGATGCTTTGACCACT
ACTACTTTGCAAGCAATCTCAGGTGCTGGTTTCTCCCCAGGTGTACCAGGTATTGATATT
CTAGACAATATTATTCCATACATTGGTGGTGAAGAAGACAAGATGGAATGGGAGACCAAG
AAAATCTTGGCTCCATTAGCAGAAGACAAGACACACGTCAAACTATTGACTCCAGAAGAA
ATCAAAGTCTCTGCTCAATGTAACAGAGTCGCTGTTTCCGATGGGCACACCGAATGTATC
TCTTTGAGGTTCAAGAACAGACCTGCTCCATCCGTCGAGCAAGTCAAGACATGCCTAAAA
GAATACGTCTGCGATGCCTACAAATTAGGCTGTCATTCTGCTCCAAAGCAAACTATTCAT
GTTTTGGAACAACCAGACAGACCTCAACCAAGGTTGGACAGGAACAGAGACAGCGGTTAC
GGTGTTTCCGTTGGTAGAATCAGAGAAGACCCATTGTTAGATTTCAAAATGGTTGTCCTT
TCCCACAACACCATTATTGGTGCCGCTGGTTCTGGTGTCTTGATTGCCGAAATCTTACTA
GCAAGAAACTTGATTTAA

>single_cluster2_YDR158W (used to calculate $\tau_{1,i}$)
ATGGCTGGAAAGAAAATTGCTGGTGTTTTGGGTGCTACTGGTTCCGTTGGTCAACGTTTC
ATTCTGTTGTTGGCAAATCACCCTCATTTCGAACTGAAAGTTCTTGGTGCCTCTTCTAGA
TCAGCTGGCAAGAAATACGTTGACGCTGTGAACTGGAAGCAAACCGATTTGCTACCGGAA
TCTGCTACCGATATTATTGTTTCCGAATGTAAATCTGAATTCTTTAAAGAGTGTGACATC
GTCTTTTCCGGATTGGATGCTGACTATGCTGGCGCTATCGAAAAGGAATTCATGGAAGCT
GGTATCGCCATTGTTTCCAATGCCAAGAATTATAGAAGAGAACAAGATGTGCCATTGATT
GTTCCTGTTGTCAATCCTGAGCATTTGGATATTGTAGCTCAAAAGCTTGACACCGCCAAG
GCTCAAGGTAAGCCAAGACCAGGGTTCATTATCTGTATTTCCAATTGTTCCACTGCAGGT
TTGGTTGCACCATTGAAGCCTTTGATTGAAAAATTCGGTCCTATTGATGCTTTGACCACT
ACTACTTTGCAAGCAATCTCAGGTGCTGGTTTCTCCCCAGGTGTACCAGGTATTGATATT
CTAGACAATATTATTCCATACATTGGTGGTGAAGAAGACAAGATGGAATGGGAGACCAAG
AAAATCTTGGCTCCATTAGCAGAAGACAAGACACACGTCAAACTATTGACTCCAGAAGAA
ATCAAAGTCTCTGCTCAATGTAACAGAGTCGCTGTTTCCGATGGGCACACCGAATGTATC
TCTTTGAGGTTCAAGAACAGACCTGCTCCATCCGTCGAGCAAGTCAAGACATGCCTAAAA
GAATACGTCTGCGATGCCTACAAATTAGGCTGTCATTCTGCTCCAAAGCAAACTATTCAT
GTTTTGGAACAACCAGACAGACCTCAACCAAGGTTGGACAGGAACAGAGACAGCGGTTAC
GGTGTTTCCGTTGGTAGGATAAGGGAGGATCCCCTTCTTGATTTCAAAATGGTTGTCCTT
TCCCACAACACCATTATTGGTGCCGCTGGTTCTGGTGTCTTGATTGCCGAAATCTTACTA
GCAAGAAACTTGATTTAA

>single_cluster3_YDR158W (used to calculate $\tau_1$)
ATGGCTGGAAAGAAAATTGCTGGTGTTTTGGGTGCTACTGGTTCCGTTGGTCAACGTTTC
ATTCTGTTGTTGGCAAATCACCCTCATTTCGAACTGAAAGTTCTTGGTGCCTCTTCTAGA
TCAGCTGGCAAGAAATACGTTGACGCTGTGAACTGGAAGCAAACCGATTTGCTACCGGAA
TCTGCTACCGATATTATTGTTTCCGAATGTAAATCTGAATTCTTTAAAGAGTGTGACATC
GTCTTTTCCGGATTGGATGCTGACTATGCTGGCGCTATCGAAAAGGAATTCATGGAAGCT
GGTATCGCCATTGTTTCCAATGCCAAGAATTATAGAAGAGAACAAGATGTGCCATTGATT
GTTCCTGTTGTCAATCCTGAGCATTTGGATATTGTAGCTCAAAAGCTTGACACCGCCAAG
GCTCAAGGTAAGCCAAGACCAGGGTTCATTATCTGTATTTCCAATTGTTCCACTGCAGGT
TTGGTTGCACCATTGAAGCCTTTGATTGAAAAATTCGGTCCTATTGATGCTTTGACCACT
ACTACTTTGCAAGCAATCTCAGGTGCTGGTTTCTCCCCAGGTGTACCAGGTATTGATATT
CTAGACAATATTATTCCATACATTGGTGGTGAAGAAGACAAGATGGAATGGGAGACCAAG
AAAATCTTGGCTCCATTAGCAGAAGACAAGACACACGTCAAACTATTGACTCCAGAAGAA
ATCAAAGTCTCTGCTCAATGTAACAGAGTCGCTGTTTCCGATGGGCACACCGAATGTATC
TCTTTGAGGTTCAAGAACAGACCTGCTCCATCCGTCGAGCAAGTCAAGACATGCCTAAAA
GAATACGTCTGCGATGCCTACAAATTAGGCTGTCATTCTGCTCCAAAGCAAACTATTCAT
GTTTTGGAACAACCAGACAGACCTCAACCAAGGTTGGACAGGAACAGAGACAGCGGTTAC
GGTGTTTCCGTTGGTAGAATCAGAGAAGACCCATTGTTAGATTTCAAAATGGTTGTCCTT
TCCCACAACACCATTATTGGTGCCGCTGGGTCGGGGGTACTTATAGCAGAGATCTTACTA
GCAAGAAACTTGATTTAA

>double_cluster1_3_YLR109W (used to calculate $\tau_2$)
ATGTCTGACTTAGTTAACAAGAAATTCCCAGCTGGCGACTACAAATTCCAGTACATAGCA
ATATCGCAGTCGGATGCTGACAGTGAATCTTGTAAGATGCCACAAACAGTTGAATGGTCC
AAATTAATTTCTGAAAACAAGAAGGTTATCATTACCGGTGCTCCAGCTGCTTTCTCCCCA
ACCTGTACTGTCAGCCATATTCCAGGTTACATCAACTACTTGGATGAATTAGTTAAGGAA
AAGGAAGTTGACCAAGTGATCGTTGTTACTGTTGACAACCCGTTCGCTAACCAAGCGTGG
GCTAAGAGTTTAGGTGTTAAGGACACCACACACATCAAGTTTGCCTCCGACCCAGGCTGT
GCTTTCACCAAATCCATTGGTTTCGAATTAGCCGTCGGTGACGGTGTTTACTGGAGTGGT
AGATGGGCCATGGTTGTTGAAAACGGTATCGTTACTTACGCTGCCAAGGAAACCAACCCA
GGTACCGATGTGACCGTATCGTCGGTAGAGTCGGTACTTGCTCATTTGTAG

>double_cluster2_3_YLR109W (used to calculate $\tau_{2,ik}$)
ATGTCTGACTTAGTTAACAAGAAATTCCCAGCTGGCGACTACAAATTCCAATACATTGCT
ATCAGCCAAAGTGATGCTGACAGTGAATCTTGTAAGATGCCACAAACAGTTGAATGGTCC
AAATTAATTTCTGAAAACAAGAAGGTTATCATTACCGGTGCTCCAGCTGCTTTCTCCCCA
ACCTGTACTGTCAGCCATATTCCAGGTTACATCAACTACTTGGATGAATTAGTTAAGGAA
AAGGAAGTTGACCAAGTGATCGTTGTTACTGTTGACAACCCGTTCGCTAACCAAGCGTGG
GCTAAGAGTTTAGGTGTTAAGGACACCACACACATCAAGTTTGCCTCCGACCCAGGCTGT
GCTTTCACCAAATCCATTGGTTTCGAATTAGCCGTCGGTGACGGTGTTTACTGGAGTGGT
AGATGGGCCATGGTAGTAGAGAATGGGATAGTAACGTACGCTGCCAAGGAAACCAACCCA
GGTACCGATGTGACCGTATCGTCGGTAGAGTCGGTACTTGCTCATTTGTAG

>wildtype_YLR109W (used to calculate $\tau_0$)
ATGTCTGACTTAGTTAACAAGAAATTCCCAGCTGGCGACTACAAATTCCAATACATTGCT
ATCAGCCAAAGTGATGCTGACAGTGAATCTTGTAAGATGCCACAAACAGTTGAATGGTCC
AAATTAATTTCTGAAAACAAGAAGGTTATCATTACCGGTGCTCCAGCTGCTTTCTCCCCA
ACCTGTACTGTCAGCCATATTCCAGGTTACATCAACTACTTGGATGAATTAGTTAAGGAA
AAGGAAGTTGACCAAGTGATCGTTGTTACTGTTGACAACCCGTTCGCTAACCAAGCGTGG
GCTAAGAGTTTAGGTGTTAAGGACACCACACACATCAAGTTTGCCTCCGACCCAGGCTGT
GCTTTCACCAAATCCATTGGTTTCGAATTAGCCGTCGGTGACGGTGTTTACTGGAGTGGT
AGATGGGCCATGGTTGTTGAAAACGGTATCGTTACTTACGCTGCCAAGGAAACCAACCCA
GGTACCGATGTGACCGTTTCCTCAGTCGAAAGTGTCTTGGCTCATTTGTAG

>single_cluster2_YLR109W (used to calculate $\tau_{1,i}$)
ATGTCTGACTTAGTTAACAAGAAATTCCCAGCTGGCGACTACAAATTCCAATACATTGCT
ATCAGCCAAAGTGATGCTGACAGTGAATCTTGTAAGATGCCACAAACAGTTGAATGGTCC
AAATTAATTTCTGAAAACAAGAAGGTTATCATTACCGGTGCTCCAGCTGCTTTCTCCCCA
ACCTGTACTGTCAGCCATATTCCAGGTTACATCAACTACTTGGATGAATTAGTTAAGGAA
AAGGAAGTTGACCAAGTGATCGTTGTTACTGTTGACAACCCGTTCGCTAACCAAGCGTGG
GCTAAGAGTTTAGGTGTTAAGGACACCACACACATCAAGTTTGCCTCCGACCCAGGCTGT
GCTTTCACCAAATCCATTGGTTTCGAATTAGCCGTCGGTGACGGTGTTTACTGGAGTGGT
AGATGGGCCATGGTAGTAGAGAATGGGATAGTAACGTACGCTGCCAAGGAAACCAACCCA
GGTACCGATGTGACCGTTTCCTCAGTCGAAAGTGTCTTGGCTCATTTGTAG

>single_cluster3_YLR109W (used to calculate $\tau_0$)
ATGTCTGACTTAGTTAACAAGAAATTCCCAGCTGGCGACTACAAATTCCAATACATTGCT
ATCAGCCAAAGTGATGCTGACAGTGAATCTTGTAAGATGCCACAAACAGTTGAATGGTCC
AAATTAATTTCTGAAAACAAGAAGGTTATCATTACCGGTGCTCCAGCTGCTTTCTCCCCA
ACCTGTACTGTCAGCCATATTCCAGGTTACATCAACTACTTGGATGAATTAGTTAAGGAA
AAGGAAGTTGACCAAGTGATCGTTGTTACTGTTGACAACCCGTTCGCTAACCAAGCGTGG
GCTAAGAGTTTAGGTGTTAAGGACACCACACACATCAAGTTTGCCTCCGACCCAGGCTGT
GCTTTCACCAAATCCATTGGTTTCGAATTAGCCGTCGGTGACGGTGTTTACTGGAGTGGT
AGATGGGCCATGGTTGTTGAAAACGGTATCGTTACTTACGCTGCCAAGGAAACCAACCCA
GGTACCGATGTGACCGTATCGTCGGTAGAGTCGGTACTTGCTCATTTGTAG

>double_cluster1_3_YPL061W (used to calculate $\tau_2$)

ATGACTAAGCTACACTTTGACACTGCTGAACCAGTCAAGATAACGCTTCCCAATGGGCTT
ACGTACGAGCAACCAACCGGTCTATTCATTAACAACAAGTTTATGAAAGCTCAAGACGGT
AAGACCTATCCCGTCGAAGATCCTTCCACTGAAAACACCGTTTGTGAGGTCTCTTCTGCC
ACCACTGAAGATGTTGAATATGCTATCGAATGTGCCGACCGTGCTTTCCACGACACTGAA
TGGGCTACCCAAGACCCAAGAGAAAGAGGCCGTCTACTAAGTAAGTTGGCTGACGAATTG
GAAAGCCAAATTGACTTGGTTTCTTCCATTGAAGCTTTGGACAATGGTAAAACTTTGGCC
TTAGCCCGTGGGGATGTTACCATTGCAATCAACTGTCTAAGAGATGCTGCTGCCTATGCC
GACAAAGTCAACGGTAGAACAATCAACACCGGTGACGGCTACATGAACTTCACCACCTTA
GAGCCAATCGGTGTCTGTGGTCAAATTATTCCATGGAACTTTCCAATAATGATGTTGGCT
TGGAAGATCGCCCCAGCATTGGCCATGGGTAACGTCTGTATCTTGAAACCCGCTGCTGTC
ACACCTTTAAATGCCCTATACTTTGCTTCTTTATGTAAGAAGGTTGGTATTCCAGCTGGT
GTCGTCAACATCGTTCCAGGTCCTGGTAGAACTGTTGGTGCTGCTTTGACCAACGACCCA
AGAATCAGAAAGCTGGCTTTTACCGGTTCTACAGAAGTCGGTAAGAGTGTTGCTGTCGAC
TCTTCTGAATCTAACTTGAAGAAAATCACTTTGGAACTAGGTGGTAAGTCCGCCCATTTG
GTCTTTGACGATGCTAACATTAAGAAGACTTTACCAAATCTAGTAAACGGTATTTTCAAG
AACGCTGGTCAAATTTGTTCCTCTGGTTCTAGAATTTACGTTCAAGAAGGTATTTACGAC
GAACTATTGGCTGCTTTCAAGGCTTACTTGGAAACCGAAATCAAAGTTGGTAATCCATTT
GACAAGGCTAACTTCCAAGGTGCTATCACTAACCGTCAACAATTCGACACAATTATGAAC
TACATCGATATCGGTAAGAAAGAAGGCGCCAAGATCTTAACTGGTGGCGAAAAAGTTGGT
GACAAGGGTTACTTCATCAGACCAACCGTTTTCTACGATGTTAATGAAGACATGAGAATT
GTTAAGGAAGAAATTTTTGGACCAGTTGTCACTGTCGCAAAGTTCAAGACTTTAGAAGAA
GGTGTCGAAATGGCTAACAGCTCTGAATTCGGTCTAGGTTCTGGTATCGAAACAGAATCT
TTGAGCACAGGTTTGAAGGTGGCCAAGATGTTGAAGGCCGGTACCGTCTGGATCAACACA
TACAACGATTTTGACTCCAGAGTTCCATTCGGTGGTGTTAAGCAATCTGGTTACGGTAGA
GAAATGGGTGAAGAAGTCTACCATGCATACACGGAGGTAAAAGCAGTAAGGATAAAGTTG
TAA

>double_cluster2_3_YPL061W (used to calculate $\tau_{2,ik}$)

ATGACTAAGCTACACTTTGACACTGCTGAACCAGTCAAGATCACACTTCCAAATGGTTTG
ACATACGAGCAACCAACCGGTCTATTCATTAACAACAAGTTTATGAAAGCTCAAGACGGT
AAGACCTATCCCGTCGAAGATCCTTCCACTGAAAACACCGTTTGTGAGGTCTCTTCTGCC
ACCACTGAAGATGTTGAATATGCTATCGAATGTGCCGACCGTGCTTTCCACGACACTGAA
TGGGCTACCCAAGACCCAAGAGAAAGAGGCCGTCTACTAAGTAAGTTGGCTGACGAATTG
GAAAGCCAAATTGACTTGGTTTCTTCCATTGAAGCTTTGGACAATGGTAAAACTTTGGCC
TTAGCCCGTGGGGATGTTACCATTGCAATCAACTGTCTAAGAGATGCTGCTGCCTATGCC
GACAAAGTCAACGGTAGAACAATCAACACCGGTGACGGCTACATGAACTTCACCACCTTA
GAGCCAATCGGTGTCTGTGGTCAAATTATTCCATGGAACTTTCCAATAATGATGTTGGCT
TGGAAGATCGCCCCAGCATTGGCCATGGGTAACGTCTGTATCTTGAAACCCGCTGCTGTC
ACACCTTTAAATGCCCTATACTTTGCTTCTTTATGTAAGAAGGTTGGTATTCCAGCTGGT
GTCGTCAACATCGTTCCAGGTCCTGGTAGAACTGTTGGTGCTGCTTTGACCAACGACCCA
AGAATCAGAAAGCTGGCTTTTACCGGTTCTACAGAAGTCGGTAAGAGTGTTGCTGTCGAC
TCTTCTGAATCTAACTTGAAGAAAATCACTTTGGAACTAGGTGGTAAGTCCGCCCATTTG
GTCTTTGACGATGCTAACATTAAGAAGACTTTACCAAATCTAGTAAACGGTATTTTCAAG
AACGCTGGTCAAATTTGTTCCTCTGGTTCTAGAATTTACGTTCAAGAAGGTATTTACGAC
GAACTATTGGCTGCTTTCAAGGCTTACTTGGAAACCGAAATCAAAGTTGGTAATCCATTT
GACAAGGCTAACTTCCAAGGTGCTATCACTAACCGTCAACAATTCGACACAATTATGAAC
TACATCGATATCGGTAAGAAAGAAGGCGCCAAGATCTTAACTGGTGGCGAAAAAGTTGGT
GACAAGGGTTACTTCATCAGACCAACCGTTTTCTACGATGTTAATGAAGACATGAGAATT
GTTAAGGAAGAAATTTTTGGACCAGTTGTCACTGTCGCAAAGTTCAAGACTTTAGAAGAA
GGTGTCGAAATGGCTAACAGCTCTGAATTCGGTCTAGGTTCTGGTATCGAAACAGAATCT
TTGAGCACAGGTTTGAAGGTGGCCAAGATGTTGAAGGCCGGTACCGTCTGGATCAACACA
TACAACGATTTTGACTCGAGGGTACCCTTCGGGGGGGTAAAGCAATCTGGTTACGGTAGA
GAAATGGGTGAAGAAGTCTACCATGCATACACGGAGGTAAAAGCAGTAAGGATAAAGTTG
TAA

>wildtype_YPL061W (used to calculate $\tau_0$)
ATGACTAAGCTACACTTTGACACTGCTGAACCAGTCAAGATCACACTTCCAAATGGTTTG
ACATACGAGCAACCAACCGGTCTATTCATTAACAACAAGTTTATGAAAGCTCAAGACGGT
AAGACCTATCCCGTCGAAGATCCTTCCACTGAAAACACCGTTTGTGAGGTCTCTTCTGCC
ACCACTGAAGATGTTGAATATGCTATCGAATGTGCCGACCGTGCTTTCCACGACACTGAA
TGGGCTACCCAAGACCCAAGAGAAAGAGGCCGTCTACTAAGTAAGTTGGCTGACGAATTG
GAAAGCCAAATTGACTTGGTTTCTTCCATTGAAGCTTTGGACAATGGTAAAACTTTGGCC
TTAGCCCGTGGGGATGTTACCATTGCAATCAACTGTCTAAGAGATGCTGCTGCCTATGCC
GACAAAGTCAACGGTAGAACAATCAACACCGGTGACGGCTACATGAACTTCACCACCTTA
GAGCCAATCGGTGTCTGTGGTCAAATTATTCCATGGAACTTTCCAATAATGATGTTGGCT
TGGAAGATCGCCCCAGCATTGGCCATGGGTAACGTCTGTATCTTGAAACCCGCTGCTGTC
ACACCTTTAAATGCCCTATACTTTGCTTCTTTATGTAAGAAGGTTGGTATTCCAGCTGGT
GTCGTCAACATCGTTCCAGGTCCTGGTAGAACTGTTGGTGCTGCTTTGACCAACGACCCA
AGAATCAGAAAGCTGGCTTTTACCGGTTCTACAGAAGTCGGTAAGAGTGTTGCTGTCGAC
TCTTCTGAATCTAACTTGAAGAAAATCACTTTGGAACTAGGTGGTAAGTCCGCCCATTTG
GTCTTTGACGATGCTAACATTAAGAAGACTTTACCAAATCTAGTAAACGGTATTTTCAAG
AACGCTGGTCAAATTTGTTCCTCTGGTTCTAGAATTTACGTTCAAGAAGGTATTTACGAC
GAACTATTGGCTGCTTTCAAGGCTTACTTGGAAACCGAAATCAAAGTTGGTAATCCATTT
GACAAGGCTAACTTCCAAGGTGCTATCACTAACCGTCAACAATTCGACACAATTATGAAC
TACATCGATATCGGTAAGAAAGAAGGCGCCAAGATCTTAACTGGTGGCGAAAAAGTTGGT
GACAAGGGTTACTTCATCAGACCAACCGTTTTCTACGATGTTAATGAAGACATGAGAATT
GTTAAGGAAGAAATTTTTGGACCAGTTGTCACTGTCGCAAAGTTCAAGACTTTAGAAGAA
GGTGTCGAAATGGCTAACAGCTCTGAATTCGGTCTAGGTTCTGGTATCGAAACAGAATCT
TTGAGCACAGGTTTGAAGGTGGCCAAGATGTTGAAGGCCGGTACCGTCTGGATCAACACA
TACAACGATTTTGACTCCAGAGTTCCATTCGGTGGTGTTAAGCAATCTGGTTACGGTAGA
GAAATGGGTGAAGAAGTCTACCATGCATACACTGAAGTAAAAGCTGTCAGAATTAAGTTG
TAA

>single_cluster2_YPL061W (used to calculate $\tau_{1,i}$)

ATGACTAAGCTACACTTTGACACTGCTGAACCAGTCAAGATCACACTTCCAAATGGTTTG
ACATACGAGCAACCAACCGGTCTATTCATTAACAACAAGTTTATGAAAGCTCAAGACGGT
AAGACCTATCCCGTCGAAGATCCTTCCACTGAAAACACCGTTTGTGAGGTCTCTTCTGCC
ACCACTGAAGATGTTGAATATGCTATCGAATGTGCCGACCGTGCTTTCCACGACACTGAA
TGGGCTACCCAAGACCCAAGAGAAAGAGGCCGTCTACTAAGTAAGTTGGCTGACGAATTG
GAAAGCCAAATTGACTTGGTTTCTTCCATTGAAGCTTTGGACAATGGTAAAACTTTGGCC
TTAGCCCGTGGGGATGTTACCATTGCAATCAACTGTCTAAGAGATGCTGCTGCCTATGCC
GACAAAGTCAACGGTAGAACAATCAACACCGGTGACGGCTACATGAACTTCACCACCTTA
GAGCCAATCGGTGTCTGTGGTCAAATTATTCCATGGAACTTTCCAATAATGATGTTGGCT
TGGAAGATCGCCCCAGCATTGGCCATGGGTAACGTCTGTATCTTGAAACCCGCTGCTGTC
ACACCTTTAAATGCCCTATACTTTGCTTCTTTATGTAAGAAGGTTGGTATTCCAGCTGGT
GTCGTCAACATCGTTCCAGGTCCTGGTAGAACTGTTGGTGCTGCTTTGACCAACGACCCA
AGAATCAGAAAGCTGGCTTTTACCGGTTCTACAGAAGTCGGTAAGAGTGTTGCTGTCGAC
TCTTCTGAATCTAACTTGAAGAAAATCACTTTGGAACTAGGTGGTAAGTCCGCCCATTTG
GTCTTTGACGATGCTAACATTAAGAAGACTTTACCAAATCTAGTAAACGGTATTTTCAAG
AACGCTGGTCAAATTTGTTCCTCTGGTTCTAGAATTTACGTTCAAGAAGGTATTTACGAC
GAACTATTGGCTGCTTTCAAGGCTTACTTGGAAACCGAAATCAAAGTTGGTAATCCATTT
GACAAGGCTAACTTCCAAGGTGCTATCACTAACCGTCAACAATTCGACACAATTATGAAC
TACATCGATATCGGTAAGAAAGAAGGCGCCAAGATCTTAACTGGTGGCGAAAAAGTTGGT
GACAAGGGTTACTTCATCAGACCAACCGTTTTCTACGATGTTAATGAAGACATGAGAATT
GTTAAGGAAGAAATTTTTGGACCAGTTGTCACTGTCGCAAAGTTCAAGACTTTAGAAGAA
GGTGTCGAAATGGCTAACAGCTCTGAATTCGGTCTAGGTTCTGGTATCGAAACAGAATCT
TTGAGCACAGGTTTGAAGGTGGCCAAGATGTTGAAGGCCGGTACCGTCTGGATCAACACA
TACAACGATTTTGACTCGAGGGTACCCTTCGGGGGGGTAAAGCAATCTGGTTACGGTAGA
GAAATGGGTGAAGAAGTCTACCATGCATACACTGAAGTAAAAGCTGTCAGAATTAAGTTG
TAA

>single_cluster3_YPL061W (used to calculate $\tau_1$)
ATGACTAAGCTACACTTTGACACTGCTGAACCAGTCAAGATCACACTTCCAAATGGTTTG
ACATACGAGCAACCAACCGGTCTATTCATTAACAACAAGTTTATGAAAGCTCAAGACGGT
AAGACCTATCCCGTCGAAGATCCTTCCACTGAAAACACCGTTTGTGAGGTCTCTTCTGCC
ACCACTGAAGATGTTGAATATGCTATCGAATGTGCCGACCGTGCTTTCCACGACACTGAA
TGGGCTACCCAAGACCCAAGAGAAAGAGGCCGTCTACTAAGTAAGTTGGCTGACGAATTG
GAAAGCCAAATTGACTTGGTTTCTTCCATTGAAGCTTTGGACAATGGTAAAACTTTGGCC
TTAGCCCGTGGGGATGTTACCATTGCAATCAACTGTCTAAGAGATGCTGCTGCCTATGCC
GACAAAGTCAACGGTAGAACAATCAACACCGGTGACGGCTACATGAACTTCACCACCTTA
GAGCCAATCGGTGTCTGTGGTCAAATTATTCCATGGAACTTTCCAATAATGATGTTGGCT
TGGAAGATCGCCCCAGCATTGGCCATGGGTAACGTCTGTATCTTGAAACCCGCTGCTGTC
ACACCTTTAAATGCCCTATACTTTGCTTCTTTATGTAAGAAGGTTGGTATTCCAGCTGGT
GTCGTCAACATCGTTCCAGGTCCTGGTAGAACTGTTGGTGCTGCTTTGACCAACGACCCA
AGAATCAGAAAGCTGGCTTTTACCGGTTCTACAGAAGTCGGTAAGAGTGTTGCTGTCGAC
TCTTCTGAATCTAACTTGAAGAAAATCACTTTGGAACTAGGTGGTAAGTCCGCCCATTTG
GTCTTTGACGATGCTAACATTAAGAAGACTTTACCAAATCTAGTAAACGGTATTTTCAAG
AACGCTGGTCAAATTTGTTCCTCTGGTTCTAGAATTTACGTTCAAGAAGGTATTTACGAC
GAACTATTGGCTGCTTTCAAGGCTTACTTGGAAACCGAAATCAAAGTTGGTAATCCATTT
GACAAGGCTAACTTCCAAGGTGCTATCACTAACCGTCAACAATTCGACACAATTATGAAC
TACATCGATATCGGTAAGAAAGAAGGCGCCAAGATCTTAACTGGTGGCGAAAAAGTTGGT
GACAAGGGTTACTTCATCAGACCAACCGTTTTCTACGATGTTAATGAAGACATGAGAATT
GTTAAGGAAGAAATTTTTGGACCAGTTGTCACTGTCGCAAAGTTCAAGACTTTAGAAGAA
GGTGTCGAAATGGCTAACAGCTCTGAATTCGGTCTAGGTTCTGGTATCGAAACAGAATCT
TTGAGCACAGGTTTGAAGGTGGCCAAGATGTTGAAGGCCGGTACCGTCTGGATCAACACA
TACAACGATTTTGACTCCAGAGTTCCATTCGGTGGTGTTAAGCAATCTGGTTACGGTAGA
GAAATGGGTGAAGAAGTCTACCATGCATACACGGAGGTAAAAGCAGTAAGGATAAAGTTG
TAA
